# Broad-spectrum antiviral activity of 3D8, a nucleic acid-hydrolyzing single chain variable fragment (scFv), targeting SARS-CoV-2 and multiple coronaviruses *in vitro*

**DOI:** 10.1101/2020.11.25.398909

**Authors:** Gunsup Lee, Shailesh Budhathoki, Geum-Young Lee, Kwang-ji Oh, Yeon Kyoung Ham, Young Jun Kim, Ye Rin Lim, Phuong Thi Hoang, Yongjun Lee, Seok-Won Lim, Jun-Mo Kim, Seungchan Cho, Tai-Hyun Kim, Jin-Won Song, Sukchan Lee, Won-Keun Kim

## Abstract

**Background:** The current pandemic, severe acute respiratory syndrome coronavirus 2 (SARS-CoV-2) is responsible for the etiology of Coronavirus-induced disease 19 (COVID-19) and poses a critical public health threat worldwide. Effective therapeutics and vaccines against multiple coronaviruses remain unavailable. Single chain variable fragment (scFv), a recombinant antibody exhibits broad-spectrum antiviral activity against DNA and RNA viruses owing to its nucleic acid-hydrolyzing property.

**Objective:** This study is aimed to investigate an antiviral activity of 3D8 scFv against SARS-CoV-2 and other coronaviruses.

**Methods:** 3D8, a recombinant scFv antibody was evaluated for antiviral activity against SARS-CoV-2, HCoV-OC43 and PEDV in Vero E6 cell cultures. Viral growth was quantified with quantitative RT-qPCR and plaque assay. Nucleic acid hydrolyzing activity of 3D8 was assessed through abzyme assays of *in vitro* viral transcripts and cell viability was determined by MTT assay.

**Results:** 3D8 inhibited the replication of SARS-CoV-2, human coronavirus OC43 (HCoV-OC43), and porcine epidemic diarrhea virus (PEDV). Our results revealed the prophylactic and therapeutic effects of 3D8 scFv against SARS-CoV-2 in Vero E6 cells. Immunoblot and plaque assays showed the reduction of coronavirus nucleoproteins and infectious particles respectively in 3D8 scFv-treated cells.

**Conclusions:** This data demonstrates the broad-spectrum antiviral activity of 3D8 against SARS-CoV-2 and other coronaviruses. Thus, it could be considered a potential antiviral countermeasure against SARS-CoV-2 and zoonotic coronaviruses.

## Introduction

Coronaviruses (subfamily Orthocoronavirinae in the family Coronaviridae of the order Nidovirales) are a group of enveloped viruses containing a single-stranded positive-sense RNA genome^1, 2^. Divergent coronaviruses constitute four genetic lineage groups, namely alphacoronaviruses, betacoronaviruses, gammacoronaviruses, and deltacoronaviruses. These viruses infect a broad range of natural reservoir hosts, including humans, bats, rodents, pigs, and camels^3^. Currently, six coronavirus species cause infectious diseases in humans. Four coronaviruses, human coronavirus OC43 (HCoV-OC43), 229E, NL63, and HKU1, induce flu-like common cold symptoms in immunocompromised individuals ^4, 5^. Two highly transmissible and pathogenic viruses, severe acute respiratory syndrome coronavirus 1 (SARS-CoV-1) and Middle East respiratory syndrome coronavirus (MERS-CoV), are associated with fatal illness involving pneumonia and respiratory disorders ^6^. In late December 2019, the city of Wuhan in the Hubei province of China, reported several cases of patients with severe pneumonia^7^, which was identified as the novel coronavirus disease (COVID-19) caused by SARS-CoV-2 ^8^. SARS-CoV-2 has several similarities to SARS-CoV and binds to the common host cell receptor, angiotensin-converting enzyme 2 (ACE-2), and Transmembrane Serine Protease 2 (TMPRSS2) ^9, 10^; however, the novel strain is genetically distinct from SARS-CoV-1^11^. The novel coronavirus SARS-CoV-2 is transmitted through species barriers from bats to humans^12^. COVID-19 is characterized by influenza-like symptoms ranging from mild to severe lung injury and multi-organ failure, leading to death in patients with comorbidities ^13^. This novel virus has led to a global pandemic, which resulted in unparalleled public health emergencies ^14^. As of February 8^th^, 2021, it has rapidly spread to 223 countries and territories, infecting over 105.8 million people including more than 2.3 million deaths ^15^.

The rapid and widespread emergence of SARS-CoV-2 presents the urgent need for antiviral countermeasures. Currently, there are no available therapeutics against human coronaviruses. Various antivirals are currently being repositioned using clinical trials: Nucleoside analogues (remdesivir, favipiravir and ribavirin), protease inhibitors (disulfiram, lopinavir and ritonavir), antiparasitic drugs (chloroquine and hydrochloroquine), pegylated interferons, monoclonal antibodies, oligonucleotide-based therapies, peptides and small-molecule drugs are contemplated for possible therapeutic agents against SARS-CoV-2 ^16, 17^. In particular, remdesivir, which inhibits viral RNA polymerases, is proposed as a potent antiviral against SARS-CoV-2 and clinical improvements have been observed in patients under compassionate use. ^5, 18^

3D8, a 27-kDa recombinant antibody fragment, is a single-chain variable fragment (scFv) that comprises a variable region of a heavy chain covalently linked to the corresponding variable region of a light chain. The protein was originally found in autoimmune-prone Murphy Roths Large (MRL) mice ^19^. The 3D8 scFv, which possesses the nucleic acid hydrolyzing activity, degrades viral DNA and/or mRNA in the infected cells ^20, 21^. This protein elicited broad-spectrum antiviral effects against classical swine fever virus^21^, influenza virus^22^, herpes simplex virus (HSV) and pseudorabies virus (PRV)^23^ *in vitro*. 3D8 exhibited *in vivo* antiviral therapeutic effects against PRV in C57BL/6 mice^24^. The transmission of avian influenza and bronchitis viruses was suppressed in transgenic chickens expressing 3D8 scFv ^25, 26^. However, the antiviral activity of 3D8 scFv against SARS-CoV-2 and other coronaviruses remains unknown.

Here, this study aimed to investigate the antiviral activity of 3D8 scFv against emerging coronaviruses *in vitro*. These data provide insight into a broad-spectrum antiviral agent against SARS-CoV-2 and multiple coronaviruses.

## Materials and Methods

### Ethics

Antiviral study of 3D8 scFv against SARS-CoV-2 was performed at the Biosafety Level-3 (BL-3) in the Hallym Clinical and Translational Science Institute, Hallym University, Chuncheon, South Korea under guidelines and protocols approved by institutional biosafety requirements. Experiments involving HCoV-OC43 and PEDV were performed in Biosafety Level-2.

### Cells and viruses

African green monkey kidney epithelial Vero cells (ATCC® CCL-81) and Vero E6 cells (ATCC® CRL-1586) were maintained in Dulbecco’s modified Eagle’s medium (12-604F, Lonza, BioWhittaker^®^, Walkersville, MD, USA) supplemented with 10% Fetal bovine serum (FBS, Gibco, Life Technologies, Grand Island, NY, USA), 1% 10 mM HEPES in 0.85% NaCl (17-737E, Lonza, BioWhittaker^®^, Walkersville, MD, USA) and 100 U/ml Penicillin-100 µg/ml Streptomycin (15140-122, Gibco, Life Technologies, Grand Island, NY, USA). Cell cultures were maintained at 37 °C with 5% CO_2_ atmosphere. SARS-CoV-2 (NCCP No. 43326) was obtained from the National Culture Collection for Pathogens (Osong, Republic of Korea). HCoV-OC43 (KBPV-VR-8) and PEDV (CV777) were obtained from Korea Bank for Pathogenic Viruses (Seoul, Republic of Korea) and the Korean Animal and Plant Quarantine Agency (Kimcheon, Republic of Korea), respectively. The viruses were propagated in Vero E6 cells, and the infectious virus titer was determined using plaque assays.

### *In vitro* transcription

To determine whether the virus gene is directly degraded by 3D8, virus genes were amplified using PCR, and *in vitro* transcription and abzyme assays were performed. cDNA was synthesized from 0.5 µg total RNA using random hexamers and MMLV reverse transcriptase (SuperScript IV Reverse Transcriptase, Thermo Fisher Scientific, Life Technologies, Carlsbad, CA, USA). PCR amplifications were performed in 20 µL reaction mixtures containing 10 µL 2× PCR premix (K-2018-1, AccuPower®, Bioneer, Oakland, USA), 1 µL forward primer, 1 µL reverse primer, and 1 µL total cDNA. All virus-specific primers were designed using the Primer 3 program (**Table S1**). The PCR program was initiated with one cycle at 95 °C for 2 min, followed by 30 cycles at 95 °C for 30 s, 45 °C for 30 s, and 72 °C for 90 s, and a final cycle at 72 °C for 5 min. Virus -specific PCR products were cloned into pGEM-T Easy Vector (A1360, Promega Co., Madison, WI, USA) and used to transform E. coli DH5a cells. In vitro transcription was reacted for 2 h at 37 °C using 2 µL T7 RNA polymerase mix, 10 mM of each dNTP, 1 µg template, and 2 µL 10× buffer. The synthesized viral RNA fragments were reacted with 3D8 and 1x reaction buffer (1× TBS, 10 mM MgCl_2_) and confirmed using electrophoresis.

### Plaque assay

Vero E6 cells were seeded at 1×10^6^ cells per well in 6 well plate (Falcon™ 353046, Corning® NY, USA) and cultured at 37 °C in CO_2_ incubator. Cells were washed with phosphate buffer saline (PBS, Lonza, USA, BioWhittaker^®^) and infected with ten-fold serial dilutions of virus suspension made in serum free maintenance media (DMEM only) and incubated at 37 °C. After the infection for 1 hour with intermittent shaking at every 15 min interval, virus inoculum was aspirated and replenished with 3 ml of over lay media (DMEM/F12 media) containing 0.6% oxoid agar and incubated at 37 °C for 4 days. The plates were then fixed with 4% Paraformaldehyde (F1119Z21 YR, Biosesang, Seongnam-si, ROK). Overlay agar media was flicked and the plates were stained with crystal violet (0.1% crystal violet in 20% methanol) for 30 min. Plaques were enumerated, and the virus titers were quantified.

### *In vitro* antiviral activity

Vero E6 cells were seeded (1×10^6^ cells per well) in 6 well plates (Corning®) and incubated overnight at 37 °C in CO_2_ incubator. The cells were washed twice with PBS, and infected with viruses at different multiplicities of infectivity (MOI) for 2 h at 37 °C. The plates were manually rocked to ensure uniform and efficient distribution of the inoculum every 15-20 minutes. After adsorption, cells were treated with 3D8 at various concentrations. In case of prophylactic treatment, Vero E6 cells were treated with 3D8 and incubated overnight before the virus challenge. Thereafter, supernatants and cells were harvested 48 h post-infection (hpi). The samples were stored at −80 °C until use.

### Real-time quantitative polymerase chain reaction

Total RNA was extracted using TRIzol (15596026, Ambion, Life Technologies, Carlsbad, CA, USA). Reverse Transcription of RNA into cDNA was performed using High Capacity RNA-to-cDNA kit (4387406, Thermo Fisher Scientific Baltics UAB, Vilnius, Lithuania) according to manufacturer’s protocol. Briefly, 1 µg of RNA was used and cDNA was synthesized by oligo deoxythymine (dT) kit. The reaction was performed at 37 °C for 60 mins followed by 95 °C for 5 mins.

Viral RNA was quantified using Power SYBR^®^ Green PCR Master Mix (4367659, Applied Biosystems™, Life Technologies Ltd., Woolston Warrington, UK) and Primers for SARS-CoV-2 and other corona viruses, with *GAPDH* as an endogenous control. Details of primers are listed in the table S2.

### Cell viability assay

Vero E6 cells were seeded (4×10^4^ cells per well) in 96-well plates (Corning®) and incubated for 24 h at 37 °C in 5% CO_2_atmosphere. 3D8 was applied at various concentrations (1 μM to 40 μM) and the cells were incubated for 48 h at 37 °C. MTT solution (Intron, 10 μL) was added to each well, and the cells were incubated at 37 °C for 3 h. Thereafter, DMSO (100 µL) was added and cellular viability was measured at 595 nm.

### Immunoblot assays

Cells were lysed using RIPA lysis buffer (SC-24948, Santa Cruz Biotechnology, Texas, USA). Cell lysates were subjected to 10% sodium dodecyl sulfate-polyacrylamide gel electrophoresis (SDS-PAGE) and transferred to a nitrocellulose membrane using a wet method. After transmembrane transfer, it was incubated with primary antibodies; anti-SARS-CoV-2 (PA1-41098, Invitrogen, Thermo Fisher Scientific, Massachusetts, USA), anti-PEDV (9191, Median Diagnostics, Chuncheon-si, ROK), anti-HCoV-OC43(LS-C79764, LS-bio, Seattle, WA, USA) and anti-human GAPDH antibody (ab9485, Abcam, Cambridge, UK) overnight at 4°C and then incubated with anti-mouse or rabbit IgG-HRP conjugate (209-035-082 or 309-035-003, Jackson ImmunoResearch, West Grove, PA, USA) for 1 h at 37 °C. The membrane reaction with ECL solution (W3652-050, GenDEPOT, DAWINBio, Hanam, ROK) was observed and confirmed by Chemiluminescence mode using ImageQuant LAS 500(GE).

### Immunocytochemistry

Vero E6 cells were seeded (1.5×10^4^ cells) in an 8-well chamber and incubated for 24 h. PEDV and HCoV-OC43 were infected at an MOI of 0.002 and 0.2 respectively for 2 h. About 5 μM (185μg/mL) purified 3D8 and 1% Pen/Strep antibiotic were added to DMEM supplemented with 10% FBS and incubated at 37 °C in 5% CO_2_ incubator. The cells were washed with PBS and fixed for 15 min in ice-cold methanol at room temperature. The cells were permeabilized with permeabilization buffer (421002, Biolegend®, San Diego, CA) for 10 min at room temperature. After blocking with 1% BSA and 0.3 M (22.52 mg/ml) glycine in PBST buffer for 1 h at room temperature, primary antibodies for detecting PEDV (mouse anti-PEDV monoclonal Ab, MBS313516, Mybio), HCoV-OC43 (mouse anti-OC43 monoclonal Ab, LS-C79764, LS-bio) and 3D8 (polyclonal rabbit IgG serum Ab) were incubated overnight at 4 °C. PEDV and HCoV-OC43 were incubated with TRITC-conjugated anti-mouse Ab (ab6786, Abcam) and 3D8 was incubated with Alexa 488-conjugated anti-rabbit Ab (ab150077, Abcam). The nuclei were stained with Hoechst (62249, Thermo Fisher Scientific Inc., Rockford, IL, USA) during the last 10 min of incubation at RT. Cells were mounted in mounting medium (H-1200 Vectashield, Vector Laboratories, Burlingame, CA, USA), and observed with a NIKON A1R (Eclipse A1Rsi and Eclipse Ti-E).

### Statistical Analysis

Statistical analyses of data were performed using Graph Pad Prism 8. Values are expressed in graph bars as mean ± SD of at least three independent experiments and a *p*-value <0.05 was considered statistically significant.

## Results

### 3D8 degraded *in vitro* RNA transcripts of SARS-CoV-2, HCoV-OC43, and PEDV

To investigate the direct nucleic acid hydrolyzing activity of scFv, *in vitro* RNA transcripts of SARS-CoV-2, HCoV-OC43, and PEDV were treated with 3D8. 3D8 degraded viral RNA transcripts in a time-dependent manner. Hydrolysis of SARS-CoV-2 RNA transcripts was evident after treatment with 3D8 for 15 min (**Figure 1A**). Complete hydrolysis of RNA transcripts was observed within 1 h compared to the control group treated with BSA. Degradation of HCoV-OC43 and PEDV RNA transcripts was observed after treatment with 3D8 (**Figure 1B and 1C**). The hydrolysis of HCoV-OC43 and PEDV transcripts was complete within 30 min. These data demonstrate the direct RNA hydrolyzing activity of 3D8 against multiple coronavirus transcripts *in vitro*.

**Figure 1.**
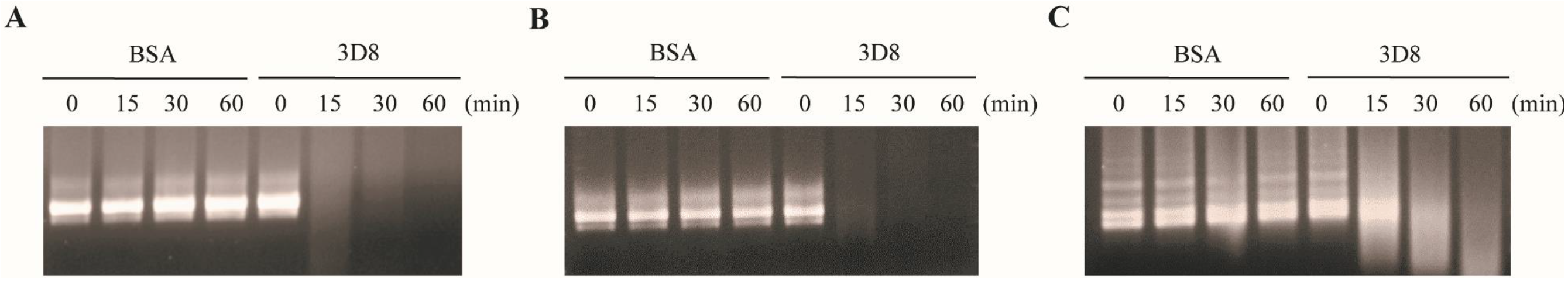
RNA hydrolyzing activity of 3D8 for *in vitro* transcripts of SARS-CoV-2, HCoV-OC43, and PEDV. (A) Nuclease activity of 3D8 against SARS-CoV-2. *In vitro* transcripts of SARS-CoV-2 were prepared and treated with 3D8. (B) Nuclease activity of 3D8 against HCoV-OC43. *In vitro* transcripts of HCoV-OC43 were prepared and treated with 3D8. (C) Nuclease activity of 3D8 against PEDV. *In vitro* transcripts of PEDV were prepared and treated with 3D8. Hydrolysis activity was determined by electrophoresis and observed under UV transilluminator.

### 3D8 inhibited SARS-CoV-2 in a dose-dependent manner

To determine the antiviral activity of 3D8 against SARS-CoV-2, various concentrations of scFv were applied to Vero E6 cells after virus challenge. SARS-CoV-2 replication in cultures treated with various doses of 3D8 was quantified using RT-qPCR **(Figure 2A)**. The replication of SARS-CoV-2 significantly decreased in a 3D8 dose-dependent manner. The 10 and 5 µM concentrations of 3D8 effectively inhibited viral replication by up to approximately 90% and 75%, respectively, compared to the non-treatment group. Infectious virus particle production was quantified using the plaque assay **(Figure 2B)**. The viral titer of SARS-CoV-2 was reduced in a 3D8 dose-dependent manner. In particular, when treated with 10 µM 3D8, the infectious particles of SARS-CoV-2 were reduced by 10 times compared to the non-treatment group. Continual treatment with 3D8 showed antiviral activity against SARS-CoV-2 at an effective concentration (EC_50_) of 4.25 µM **(Figure 2C)**. Moreover, this scFv did not show cytotoxicity in Vero E6 cells treated with 3D8 scFv at concentrations ranging from 1 µM to 10 µM **(Figure 2D)**. However, cytotoxicity was observed at 40 µM.

**Figure 2.**
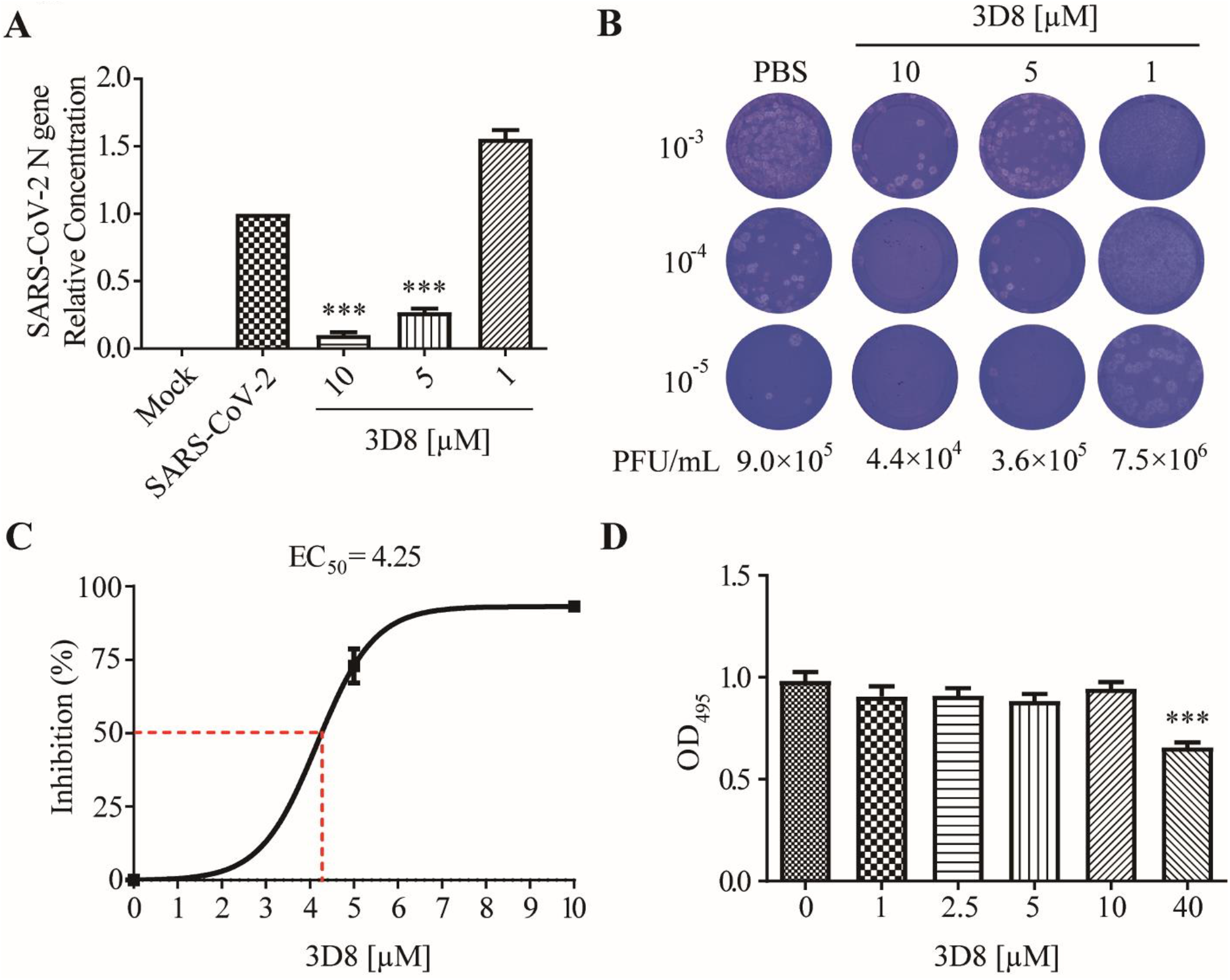
Antiviral activity of 3D8 scFv against SARS-CoV-2 in a dose-dependent manner and a cytotoxicity assay. (A) Dose-dependent inhibition of SARS-CoV-2 by 3D8. Vero E6 cells were infected with SARS-CoV-2 at 2 hpi and treated with a range of 3D8 concentrations for 48 hrs. The cells were harvested, and the viral RNA load was determined using RT-qPCR. (B) Supernatants from the 3D8-treated samples were collected, and a plaque assay was performed to determine the infectious viral titer. (C) Percent inhibition of SARS-CoV-2 replication was shown by 3D8 in Vero E6 cells. Replication was measured via quantification of the viral RNA level. (D) Cytotoxicity testing of 3D8 in Vero E6 cells was performed by applying a range of various concentrations in uninfected cell cultures. Error bars indicate the standard deviation of triplicate measurements in a representative experiment. (***p<0.001, One-way ANOVA test; ns: non-significant)

### 3D8 effectively inhibited SARS-CoV-2 in pretreated cells (prophylactic effect)

We determined the prophylactic antiviral activity of 3D8 scFv against SARS-CoV-2 in pretreated cell cultures. A significant reduction in the SARS-CoV-2 N gene copies was observed upon treatment with 10 µM 3D8 scFv **(Figure 3A)**. The N protein of SARS-CoV-2 was not observed after treatment with 3D8 scFv **(Figure 3B)**. Furthermore, we determined the inhibitory effect of 3D8 on the production of SARS-CoV-2 infectious particles. The production of infectious virus particles was more than 10 times lower in the scFv-treated group compared to the control group **(Figure 3C)**. Collectively, these data demonstrate that 3D8 has a prophylactic effect on SARS-CoV-2 infection.

**Figure 3.**
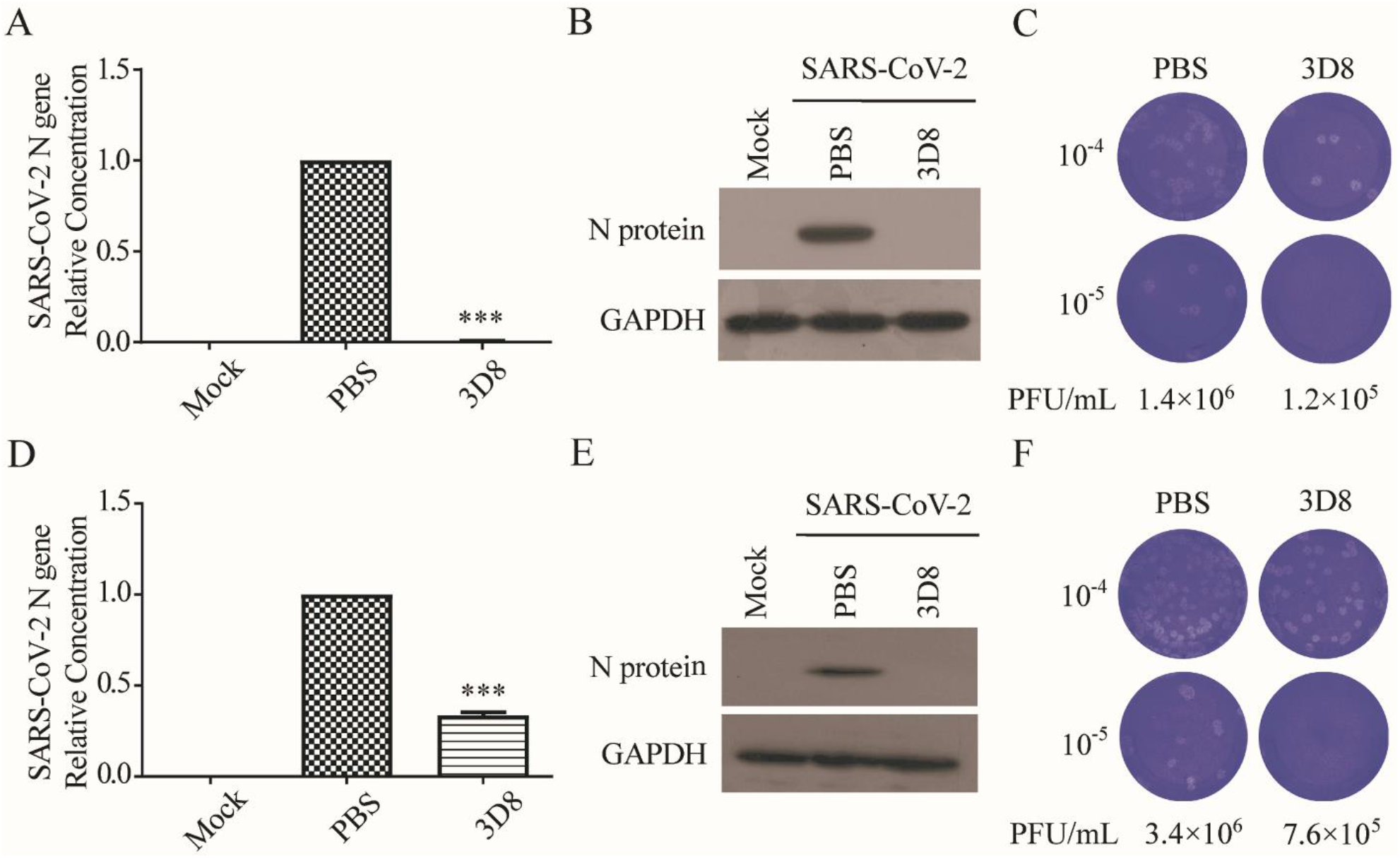
Prophylactic and therapeutic antiviral effects of 3D8 against SARS-CoV-2. (A) Inhibition of SARS-CoV-2 by 3D8 in pretreated cell cultures. Vero E6 cells pretreated with 3D8 were infected with SARS-CoV-2. At 48 hpi, the cells were harvested, and the viral copy number was quantified based on the relative concentration of the N gene. (B) The N protein of SARS-CoV-2 was shown by western blot. (C) Supernatants were harvested from 3D8-pretreated cell cultures infected with SARS-CoV-2, and the infectious viral titer was determined using a plaque assay. The clear zone indicates the plaques formed. (D) Inhibition of SARS-CoV-2 by 3D8 in post-treated cell cultures. Vero E6 cells were infected with SARS-CoV-2 at 2 hpi and treated with 3D8 for 48 hrs. The cells were harvested, and the viral copy number was quantified based on the relative concentration of the N gene. (E) The N protein of SARS-CoV-2 was shown by western blot. (F) Supernatants were harvested from 3D8 post-treated cell cultures infected with SARS-CoV-2, and the infectious viral titer was determined using a plaque assay. (***p<0.001; ****p<0.0001, One-way ANOVA test; ns: non-significant)

### 3D8 effectively inhibited SARS-CoV-2 in post-treated cells (therapeutic effect)

To determine the therapeutic effect, we assessed the inhibitory activity of 3D8 scFv on SARS-CoV-2 strain 2 hpi **(Figure 3D)**. 3D8 scFv decreased the gene copy number of the N gene. Western blot analysis revealed the absence of N protein expression in the 3D8-treated samples **(Figure 3E)**. The production of infectious SARS-CoV-2 particles in the 3D8-treated group was 10 times lower than that in the control group **(Figure 3F)**. The reduction of viral gene copy number, N protein expression, and infectious virus particles indicated the therapeutic effect of 3D8 on SARS-CoV-2 infection.

### 3D8 possesses broad-spectrum antiviral activity against multiple coronaviruses

To explore the antiviral activity of 3D8 against other coronaviruses, viral gene copies and infectious particles were examined after HCoV-OC43 and PEDV infections. The 3D8-treated group showed effective inhibition of viral replication upon HCoV-OC43 infection. The load of HCoV-OC43 RNA was significantly reduced in a 3D8 dose-dependent manner **(Figure 4A)**. The expression of viral proteins was inhibited in the presence of 3D8 **(Figure 4B)**. 3D8 effectively inhibited the replication of HCoV-OC43 with an EC_50_ value of 1.40 µM **(Figure 4C)**. Immunohistochemistry analysis revealed a reduction in HCoV-OC43 replication upon 3D8 treatment **(Figure 4D)**.

**Figure 4.**
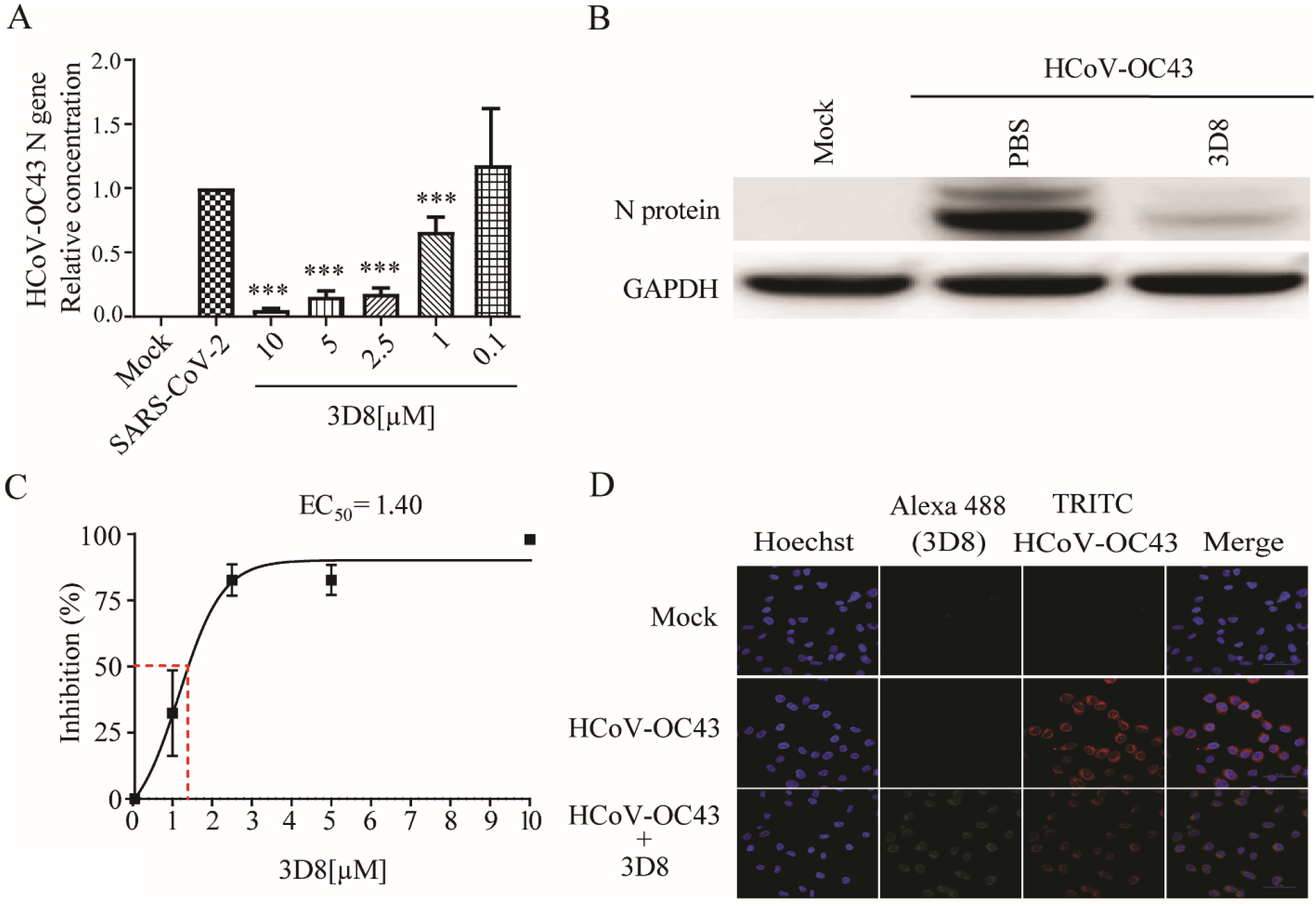
*In vitro* antiviral effect of 3D8 against human coronavirus HCoV-OC43. (A) Dose-dependent inhibition of HCoV-OC43 by 3D8. Vero E6 cells were infected with HCoV-OC43 at 2 hpi and treated with a range of 3D8 concentrations for 48 hrs. The cells were harvested, and the virus copy number was determined using qPCR. (B) The N protein of HCoV-OC43 was shown by western blot. (C) Percent inhibition of HCoV-OC43 replication was shown by 3D8 in Vero E6 cells. Replication was measured via quantification of the viral RNA level. (D) At 48 hpi, the cells were washed with PBS and fixed using methanol. Then, they were permeabilized with buffer and blocked with BSA. The cells were then incubated with primary antibodies overnight. After incubation, TRITC-conjugated anti-mouse and Alexa 488-conjugated anti-rabbit antibodies were added. Hoechst was used to stain the nucleus. (*p< 0.05; ***p<0.001, One-way ANOVA test; ns: non-significant)

Treatment with 3D8 resulted in effective inhibition of viral replication during PEDV infection. The PEDV RNA load was significantly suppressed in a 3D8 dose-dependent manner **(Figure 5A)**. The expression of viral proteins was reduced upon treatment with 3D8 **(Figure 5B)**. The EC_50_ value of 3D8 was 1.10 µM against PEDV **(Figure 5C)**. Immunohistochemistry analysis revealed a reduction in PEDV replication upon 3D8 treatment **(Figure 5D)**. These data demonstrated the broad-spectrum activity of 3D8 against multiple zoonotic coronaviruses.

**Figure 5.**
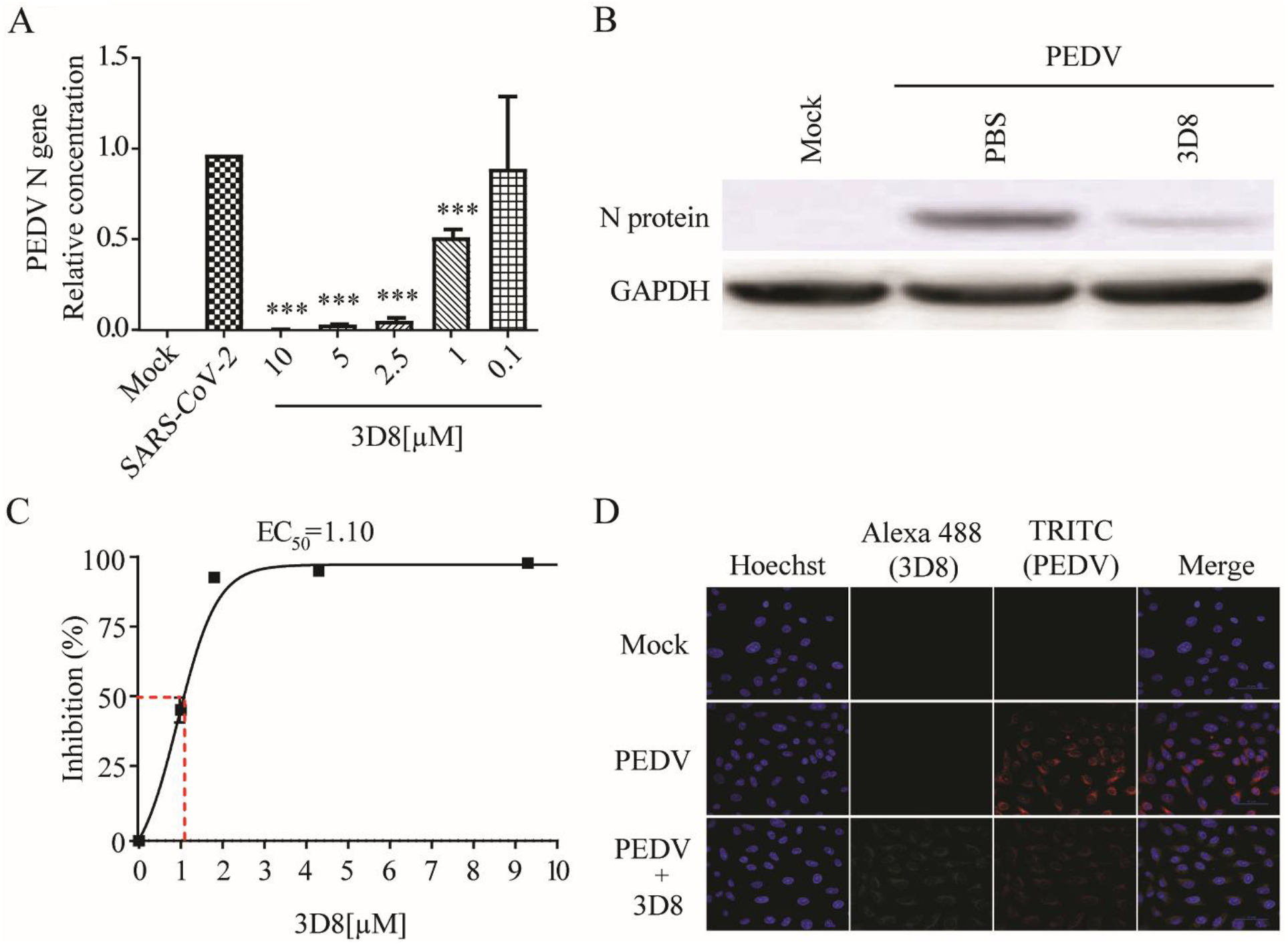
*In vitro* antiviral effect of different concentrations of 3D8 against PEDV. (A) Dose-dependent inhibition of PEDV by 3D8. Vero E6 cells were infected with PEDV at 2 hpi and treated with a range of 3D8 concentrations for 48 hrs. The cells were harvested, and the viral RNA load was determined using qPCR. (B) The N protein of PEDV was shown by western blot. (C) Percent inhibition of PEDV replication was shown by 3D8 in Vero E6 cells. Replication was measured via quantification of the viral RNA level. (D) At 48 hpi, the cells were washed with PBS and fixed with methanol. Then, they were permeabilized with buffer and blocked with BSA. The cells were incubated with primary antibodies overnight. After incubation, TRITC-conjugated anti-mouse and Alexa 488-conjugated anti-rabbit antibodies were added. Hoechst was used to stain the nucleus. (*p< 0.05; **p<0.01, One-way ANOVA test; ns: non-significant)

## Discussion

Novel human coronaviruses have emerged during the past two decades ^3, 4^. The COVID-19 outbreak occurred in late December in Wuhan (China) and rapidly became a global pandemic ^7^. The public health emergency caused by the SARS-CoV-2 outbreak presents the demand for countermeasures against emerging and re-emerging zoonotic coronaviruses. As the virus disseminates, efforts are being made to mitigate transmission via public health interventions including social distancing, case isolation (quarantine), and contact tracing. However, therapeutics and vaccines against SARS-CoV-2 are urgently needed for the effective control of outbreaks. In this study, we demonstrated that 3D8, a nucleic acid-hydrolyzing scFv, inhibited the replication of SARS-CoV-2 and multiple coronaviruses *in vitro*.

scFv is a molecule derived from an antibody composed of variable region of heavy and light chains linked with peptides^19^. This protein has applied for biotechnological and medicinal applications such as infectious diseases, cancer therapy and potential alternatives to conventional diagnostic approaches^27, 28^. The scFv has various advantages over traditional monoclonal antibodies, such as ease of genetic manipulation, rapid molecular design and characterization, greatly reduced size, production of antibodies against viral proteins, and various expression systems^28^. Neutralizing scFv against N protein protected piglets from PEDV infection. The orally administered piglets either exhibited no clinical symptoms or mild symptoms and intestinal lesions, with significantly increased survival rates^29^. 3D8, a unique scFv with broad-spectrum nuclease activity, confers antiviral activities against a variety of viruses including DNA and RNA viruses^23, 24^. 3D8 scFv showed antiviral effects against Infectious bronchitis virus, a member of gammacoronaviruses in transgenic chickens^26^. However, the antiviral activity of 3D8 scFv against SARS-CoV-2 and other coronaviruses remained to be investigated.

Our study demonstrated that 3D8 conferred effective antiviral activity against SARS-CoV-2, HCoV-OC43, and PEDV *in vitro*. 3D8 inhibited the replication of multiple coronaviruses in a dose-dependent manner. The nuclease activity of 3D8 contributed to the reduction of infectious virus particles, indicating that the degradation of viral nucleic acids prohibited the production of viral genomes and proteins **(Figure 6)**. In addition, therapeutic treatment with 3D8 scFv inhibited the replication of SARS-CoV-2, HCoV-OC43 and PEDV, resulting in the lack of viral replication, infectious particle formation and protein expression. Previous studies demonstrated the biochemical characteristics and robust antiviral activity of 3D8 scFv against Classical Swine Fever virus and HSV *in vitro*^21, 23^. Transgenic mouse and chicken were generated by expressing 3D8, demonstrating the *in vivo* antiviral activity against Influenza virus and PRV^24, 25^. The entry mechanism of 3D8 revealed a caveolin-dependent manner and easy penetration of the cell without a carrier^30^. Intranasal transfer of 3D8 scFv into a mouse penetrated well into the epithelial barrier of lung tissues^22^. 3D8 stably existed in the lung and various tissues of the inoculated mouse (not shown). Taken together, 3D8, a nucleic-acid hydrolyzing mini-antibody, may be a potential antiviral candidate due to its broad-spectrum activity, easy penetration into the cell, and the accessibility to the lung *in vivo*.

**Figure 6.**
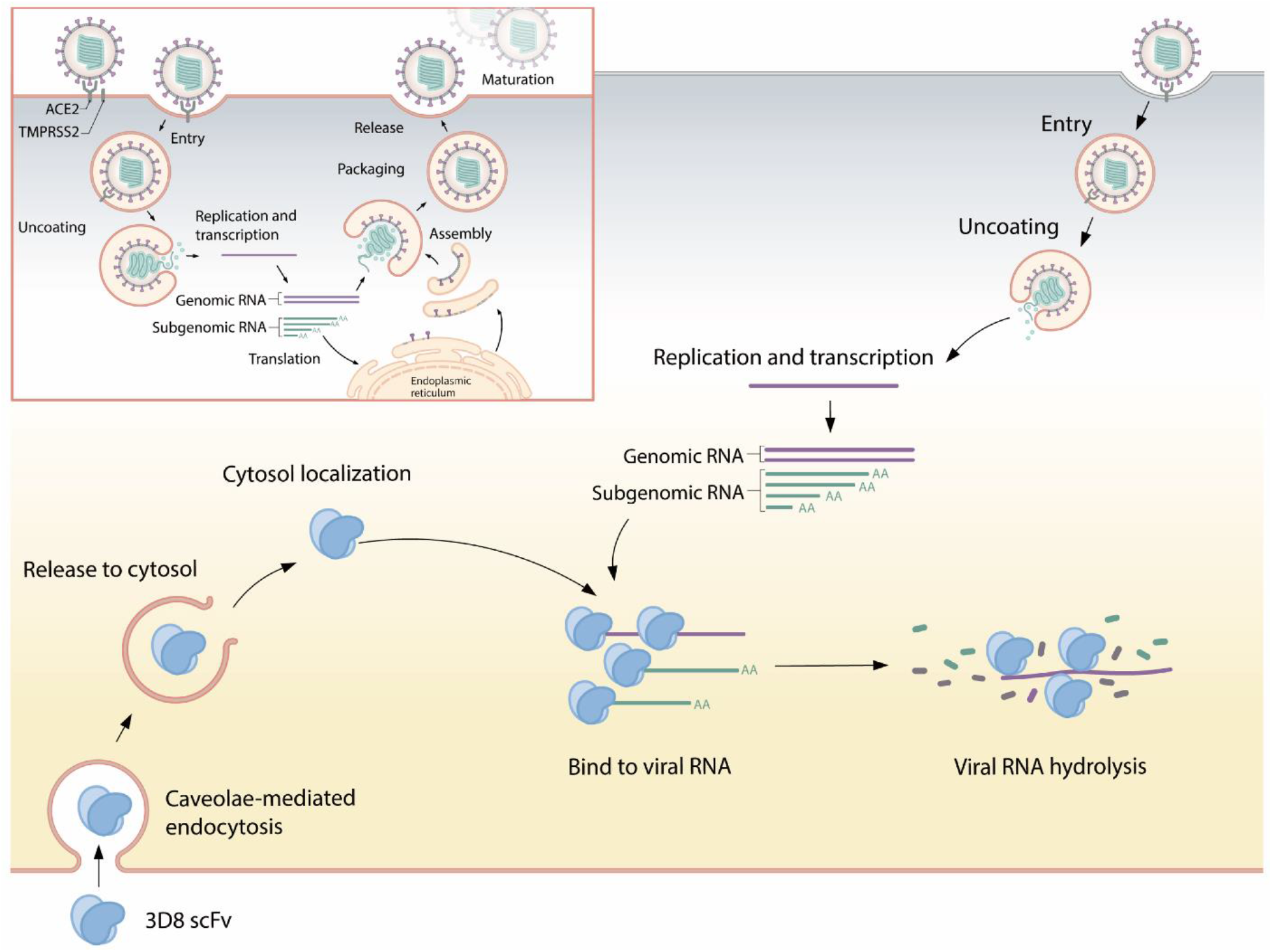
Suggested mode of action for 3D8. 3D8 scFv is internalized into the cell through caveolae-mediated endocytosis. After release from the endosomal compartment, 3D8 binds to the viral nucleic acid and degrades it to prevent its amplification, thus inhibiting viral growth. 3D8 exerts nuclease activity without sequence specificity and hydrolyzes viral RNA genomes or transcripts.

Infection of coronaviruses has significantly impacted on humans and livestocks^31^. However, the effective antiviral countermeasures against these viruses are still unavailable. HCoV-OC43, which belongs to the lineage betacoronaviruses, is associated with mild common cold in humans. HCoV-OC43 infection occurs frequently in early childhood and causes acute respiratory tract illness, pneumonia and croup^32^. PEDV, a member of alphacoronaviruses, is a highly contagious coronavirus that causes severe diarrhea and death in neonatal piglets^29^. All age groups are highly susceptible to PEDV infection, with neonatal piglets under two weeks of age showing the highest mortality rates^29^. In this study, the administration of 3D8 also elicited a reduction in replication and protein synthesis of HCoV-OC43 and PEDV, followed by the formation of infectious particles. These results suggest that 3D8 may be a potential antiviral candidate for the human public health and livestock industry.

RNA viruses constantly undergo genetic mutation owing to the lack of proofreading activity of RdRp polymerases, resulting in the emergence of new variants of the virus over time^33^. A variant of SARS-CoV-2 with D614G substitution on the spike protein was reported in February, 2020^34^. This variant quickly became a dominant strain circulating globally due to increased infectivity in humans^34, 35^. Notably, the variants of SARS-CoV-2 from the United Kingdom and the Republic of South Africa have raised significant concern^36^. The variant from United Kingdom, referred to as SARS-CoV-2 (B.1.1.7) with 23 nucleotide substitutions, was a phylogenetic genotype distinct from the origin of SARS-CoV-2 strains^37^. Another variant from South Africa, referred to as SARS-CoV-2 (B.1.351) shared a genetic similarity with SARS-CoV-2 B.1.1.7, but it belonged to a distinct phylogenetic lineage from B 1.1.7.^34, 36^ The genetic variation of SARS-CoV-2 may impact the effectiveness and efficacy of antivirals or vaccines targeting viral proteins, since genetic mutation facilitates evasion of antiviral molecular mechanism and humoral responses. In this study, *in vitro* RNA transcripts of SARS-CoV-2, HCoV-OC43, and PEDV were degraded by 3D8 scFv in a sequence-independent manner. In addition, administration of 3D8 elicited the degradation of viral RNA upon infection with several Influenza virus strains, including H1N1 and H9N2^22, 25^. Thus, 3D8 scFv possesses broad-spectrum nucleic acid hydrolyzing activity, indicating that this protein may exert antiviral activity against variants of SARS-CoV-2.

Our study focused on the inhibitory effect of 3D8 scFv against multiple coronaviruses *in vitro*, demonstrating that 3D8 effectively inhibited SARS-CoV-2 and other coronaviruses (HCoV-OC43 and PEDV). However, the absence of *in vivo* studies in animal models is a major limitation. To address this issue, further studies are needed to illustrate the *in vivo* inhibitory effect of 3D8 scFv in transgenic mice, Syrian hamsters and non-human primate models. In conclusion, 3D8 scFv confers broad-spectrum antiviral activity against SARS-CoV-2 and multiple coronaviruses. This study provides insights into the effective antiviral countermeasure of scFv against emerging viral outbreaks.

## Acknowledgment

We thank Mr. Jong Hoon Jeon (Hallym University) for supporting the BL-3 facility.

## Funding

This work was supported by Novelgen (S-2018-1158-000) and the Research Program to Solve Social Issues of the National Research Foundation of Korea (NRF) funded by the Ministry of Science and ICT (NRF-2017M3A9E4061992).

## Conflict of interest

The authors declare that the research was conducted in the absence of any commercial or financial relationships that could be construed as a potential conflict of interest.

## Supplementary Data

**Table S1.**
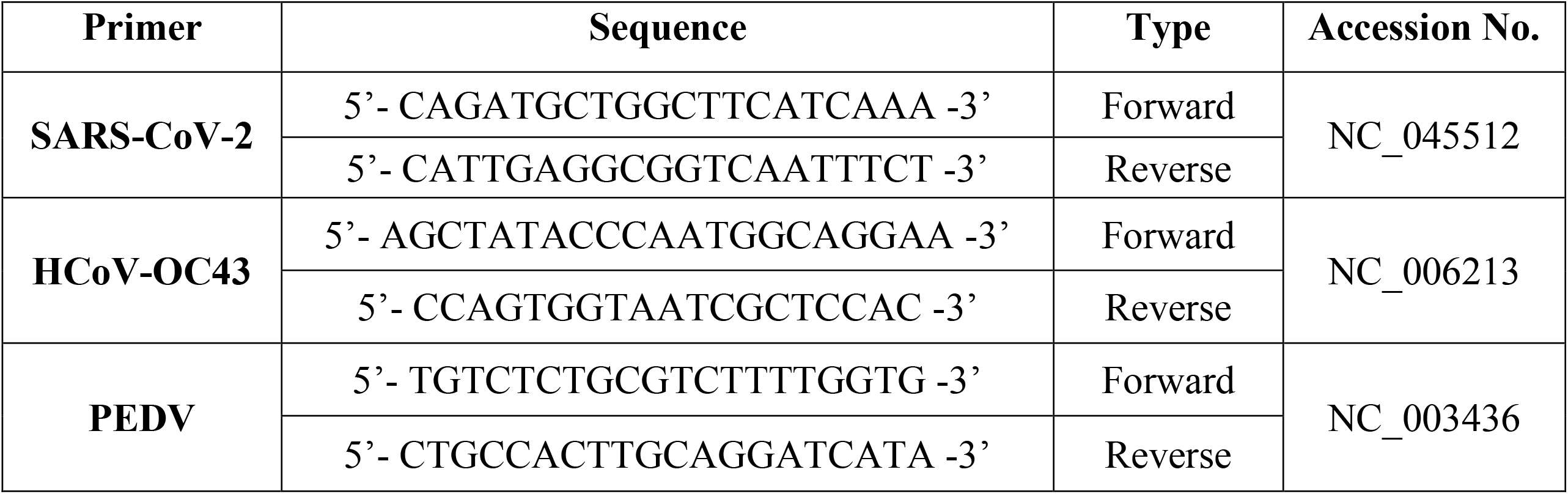
Primers used for in vitro transcription viral spike gene amplification.

**Table S2.**
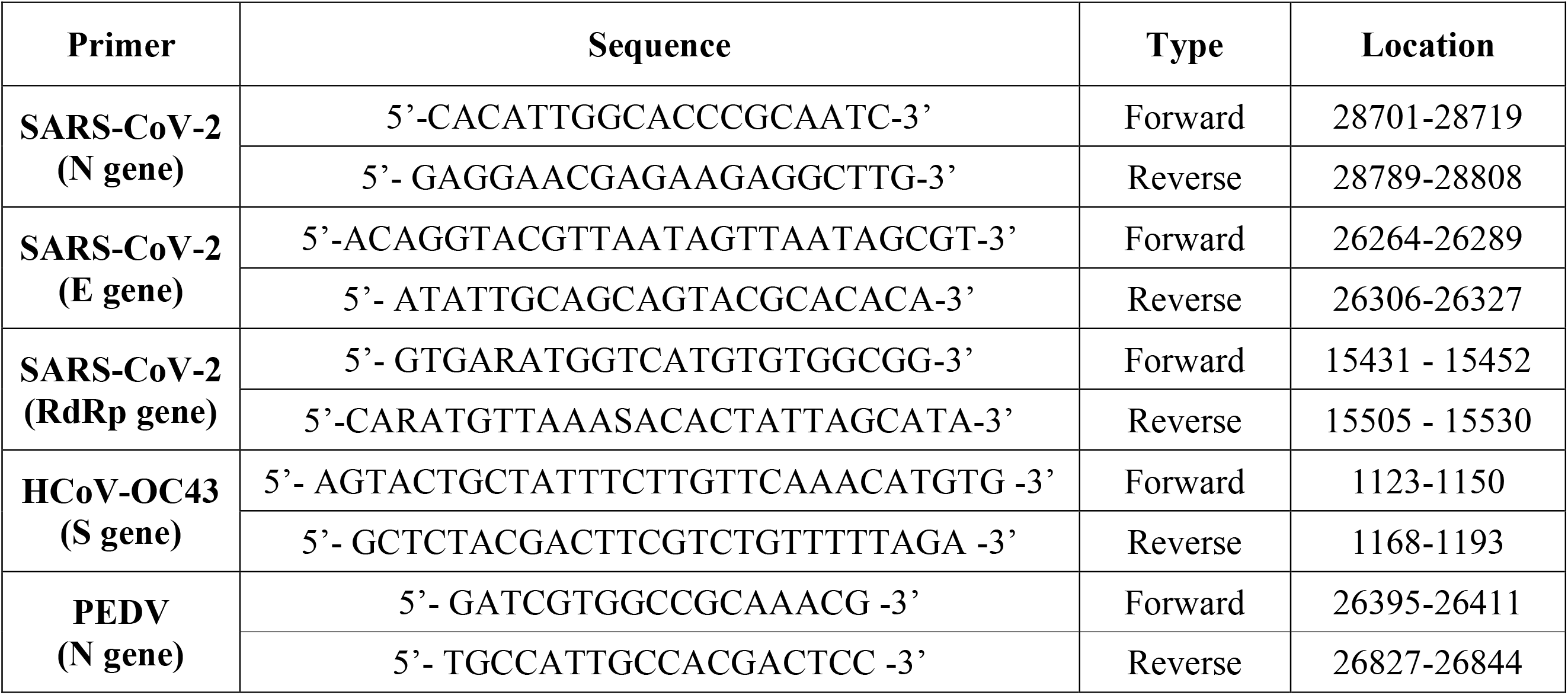
Primers used for RT-qPCR of SARS-CoV-2, HCoV-OC43, and PEDV.

## Notes

### Competing Interest Statement

The authors have declared no competing interest.

### Summary of Updates

Title changed; Main context significantly revised; Figure 1 added; All of figures updated; Supplemental tables updated.

